# Gut sulfide metabolism modulates behavior and brain bioenergetics

**DOI:** 10.1101/2025.04.09.647962

**Authors:** Roshan Kumar, Delawrence J Sykes, Victor I Band, Megan L. Schaller, Romel Patel, Victor Vitvitsky, Peter Sajjakulnukit, Rashi Singhal, Harrison K.A. Wong, Suchitra K. Hourigan, Fumito Ichinose, Costas A. Lyssiotis, Yatrik M. Shah, Ruma Banerjee

**Affiliations:** Departments of Biological Chemistry; Molecular and Integrative Physiology; Internal Medicine; Rogel Cancer Center, University of Michigan, Ann Arbor, MI, USA; Department of Biology, Berry College, Mount Berry, GA, USA; NIAID, National Institutes of Health, Bethesda, MD; Anesthesia Center for Critical Care Research, Department of Anesthesia, Critical Care and Pain Medicine, Massachusetts General Hospital and Harvard Medical School, Boston, MA, USA

**Keywords:** Hydrogen sulfide, sulfide quinone oxidoreductase, gut-brain axis, ketone body, goblet cells, ventricles, methionine rich diet

## Abstract

The host-microbiome interface is rich in metabolite exchanges and exquisitely sensitive to diet. Hydrogen sulfide (H_2_S) is present at high concentrations at this interface, and is a product of both microbial and host metabolism. The mitochondrial enzyme, sulfide quinone oxidoreductase (SQOR), couples H_2_S detoxification to oxidative phosphorylation; its inherited deficiency presents as Leigh disease. Since an estimated two thirds of systemic H_2_S metabolism originates in gut, it raises questions as to whether impaired sulfide clearance in this compartment contributes to disease, and whether it can be modulated by dietary sulfur content. In this study, we report that SQOR deficiency confined to murine intestinal epithelial cells, perturbs colon bioenergetics that is reversed by antibiotics, establishing a significant local contribution of microbial H_2_S to host physiology. We also find that a 2.5-fold higher methionine intake, mimicking the difference between animal and plant proteins, synergized with intestinal SQOR deficiency to adversely impact colon architecture and alter microbiome composition. In serum, increased thiosulfate, a biomarker of H_2_S oxidation, revealed that intestinal SQOR deficiency combined with high dietary methionine, affects sulfide metabolism globally and perturbs energy metabolism as indicated by higher ketone bodies. The mice exhibited lower exploratory locomotor activity while brain MRI revealed an atypical reduction in ventricular volume, which was associated with lower aquaporin 1 that is important for cerebrospinal fluid secretion. Our study reveals the dynamic interaction between dietary sulfur intake and sulfide metabolism at the host-microbe interface, impacting gut health, and the potential for lower dietary methionine intake to modulate pathology.

**Significance Statement:** The host-microbiome interface is rich in metabolite-based communications that are modulated by diet. Hydrogen sulfide (H_2_S), which is a respiratory poison at high concentrations, is enriched at this interface, and is detoxified by the host enzyme, sulfide quinone oxidoreductase (SQOR). Given the quantitatively significant contribution of gut to systemic H_2_S metabolism, we examined how SQOR deficiency restricted to murine intestinal epithelial cells, interacts with high dietary methionine, designed to approximate the difference between plant versus animal protein levels, to affect local and global bioenergetics. Our study revealed profound short- and long-range impacts resulting from the synergy between decreased H_2_S clearance capacity in gut and high dietary methionine on global energy metabolism, brain pathology, and behavior.

## INTRODUCTION

Historically known as an environmental toxin, hydrogen sulfide (H_2_S) is surprisingly abundant in the liminal zone between host epithelium and gut microbiota. H_2_S acts as a respiratory toxin by inhibiting complex IV in the electron transport chain (ETC), but is also a signaling molecule that mediates a plethora of physiological effects (1). The short- and long-range impacts of local host-microbiota interactions on systemic energy metabolism and organismal behavior, are however, poorly understood. Colon sulfide, which reportedly ranges from ∼0.2-2.4 mM (2, 3), is presumed to be primarily microbial in origin, generated via cysteine desulfuration or sulfate reduction reactions (4). H_2_S is also synthesized by the host enzymes, cystathionine β-synthase (5) and γ-cystathionase (6), from cysteine and/or homocysteine, both products of methionine metabolism (7), or by mercaptopyruvate sulfurtransferase, from 3-mercaptopyruvate (8, 9). Fecal sulfide levels are positively correlated with dietary intake of animal protein (10), pointing to the three-way interplay between host, microbes and diet in influencing H_2_S metabolism. The methionine content of animal protein is estimated to be ∼2.5-fold higher than in plant protein (11). When methionine is plentiful, homocysteine, a branchpoint metabolite, is directed to cysteine synthesis via the transsulfuration enzymes, cystathionine β-synthase and γ-cystathionase (12). While catalytic promiscuity of the transsulfuration enzymes contributes to H_2_S synthesis (13), their reaction specificity is regulated, e.g. by ER stress or hypoxia (14, 15).

At low concentrations, H_2_S oxidation supports ATP synthesis, funneling electrons to coenzyme Q (CoQ), while the sulfane sulfur is transferred to glutathione, in a reaction catalyzed by sulfide quinone oxidoreductase (SQOR) (16). At higher concentrations, H_2_S inhibits complex IV and increases the O_2_ pressure for half-maximal respiration (17). The metabolic memory of cellular H_2_S exposure is relatively long-lived and perturbs cellular bioenergetics (18), increasing aerobic glycolysis, glutamine-dependent reductive carboxylation, and lipid biogenesis (9, 19, 20). The other enzymes in the mitochondrial sulfide oxidation pathway include ETHE1, a persulfide dioxygenase (21), TST, a thiosulfate sulfurtransferase (22), and sulfite oxidase (23).

Inherited deficiency of SQOR presents as Leigh syndrome (24), a phenotypically heterogenous mitochondrial disease that is often characterized by diffuse multifocal spongiform degeneration in brain (25). Fasting or infection often trigger the clinical manifestations of Leigh disease, which is caused by mutations in >80 genes; a subset of these patients present with gastrointestinal problems. Two of the three SQOR deficiency patients described so far, succumbed to their illness at age 4 and 8, respectively, while the third, survived at least into adolescence, despite recurrent encephalopathic episodes (24). Involvement of the gut-brain axis in SQOR-deficiency Leigh disease is suggested by the efficacy of a sulfur restricted diet (11) or metronidazole, a broad spectrum antibiotic, to attenuate associated pathology in a murine model (26). These mice, expressing SQOR in the cytoplasm instead of mitochondria, exhibit a greatly reduced life span, albeit one that increased from ∼12 to 26 week median survival with metronidazole (26, 27). Two other whole-body SQOR knockout models also exhibited decreased lifespan with all mice dying within 7-15 weeks of age (28, 29). A sulfur-restricted diet or metronidazole also improves clinical outcomes in patients with ethylmalonic encephalopathy due to inherited deficiency of ETHE1, the second enzyme in the sulfide oxidation pathway (30, 31). Both SQOR and ETHE1 deficiency are characterized by increased H_2_S and decreased complex IV activity. The protective effect of antibiotics in the backdrop of decreased H_2_S oxidation capacity due to ETHE1 deficiency, is correlated with lower H_2_S and complex IV activity in liver, brain and muscle (32).

Since the gut microbe-host interface is estimated to account for ∼70% of systemic sulfide homeostasis (33), we posited that the etiology of sulfide metabolism disorders is connected to loss of intestinal H_2_S oxidation capacity. In the present study, Villin Cre recombinase was used to specifically dissect how SQOR deficiency in intestinal epithelial cells (IEC) interacts with dietary methionine modulation, to impact bioenergetics, brain pathology, and behavior. We demonstrate that antibiotics restore MT-CO1 levels, a sensitive indicator of complex IV and ETC capacity, and increase electron acceptor availability in mice lacking IEC SQOR. In addition to the adverse local effects of a methionine rich diet (MRD) on colon, it synergizes with IEC SQOR deficiency to exert pleiotropic effects on gut microbial composition, ketogenesis, brain pathology and animal behavior. Elevated serum thiosulfate, a stable biomarker of H_2_S oxidation, indicates that elevated circulating sulfide *per se* is culpable for the long range effects in IEC SQOR deficiency mice on MRD. Our study reveals the critical importance of SQOR as an intestinal shield against microbial sulfide for maintaining gut health that is modulated by dietary methionine, and when compromised, is associated with pathology along the gut-brain axis.

## RESULTS

### Antibiotics reverse colon bioenergetic impairments due to gut SQOR deficiency

The interplay between gut sulfide oxidation capacity and microbial sulfide production on host bioenergetics (Fig. 1A) was assessed using an IEC-specific SQOR knockout (KO) mouse, Villin^Cre^ Sqor^fl/fl^ (Supplementary Fig. 1A,B). Greatly diminished levels of MT-CO1, the copper- and heme-containing subunit of complex IV, indicated that ETC function was compromised due to SQOR KO (Fig. 1B,C). MT-CO1 loss has been seen previously in cells and tissue under chronic sulfide stress (17, 32). Decreased ETC function was correlated with a marked reductive shift in the CoQ_9_ and CoQ_10_ pools (Fig 1D) although the total CoQ pool size was unaffected (Supplementary Fig. 1C-E). In mice, CoQ_9_ and CoQ_10_ are the major and minor cofactor forms, respectively (34). Remarkably, antibiotics reversed the effect of the SQOR deficiency genotype on MT-CO1 levels (Fig 1D,E) and restored the CoQ redox status (Fig. 1G and Supplementary Fig. 1F,G). Together, these data reveal that gut H_2_S is largely derived from microbiota, and that IEC SQOR is essential for shielding the host against the toxic effects of sulfide (Fig. 1H).

**Figure 1.**
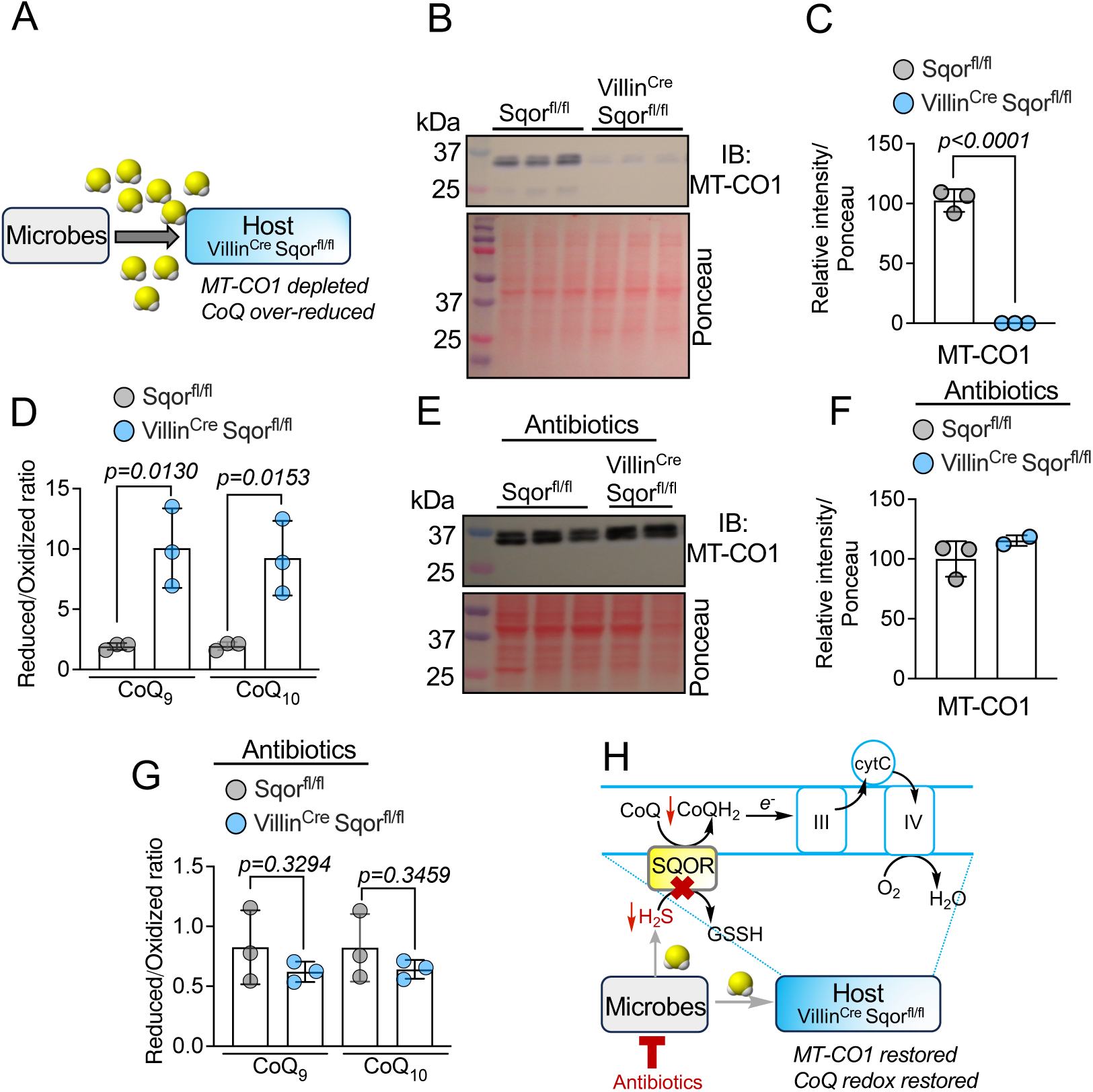
Sqor KO in IEC induces a reductive shift in the CoQ pool that is reversed by antibiotics. **(A)** Scheme showing microbiota-derived H2S lowers MT-CO1 and over-reduces the CoQ pool in Villin^Cre^ Sqor^fl/fl^ mouse colon. **(B-C)** Western blot (B) and quantitative analysis (C) of MT-CO1 in Sqor^fl/fl^ and Villin^Cre^ Sqor^fl/fl^ mice (n=3 independent mice). **(D)** Influence of SQOR deficiency on CoQ_9_ and CoQ_10_ redox poise in colon (n=3 independent mice). **(E, F)** Effect of antibiotics on colon MT-CO1 levels as detected by Western blot analysis (n=3 independent mice for Sqor^fl/fl^ and n=2 for Villin^Cre^ Sqor^fl/fl^ mice) (E) and the CoQ_9_ and CoQ_10_ redox poise (n=3 independent Sqor^fl/fl^ and Villin^Cre^ Sqor^fl/fl^ mice) (F). **(G)** Effect of antibiotics on SQOR deficiency on CoQ_9_ and CoQ_10_ redox status in colon (n=3 independent mice). **(H)** Scheme showing antibiotics decreases colon exposure to microbial H2S, which protects colon ETC function in Villin^Cre^ Sqor^fl/fl^ mice.

### High dietary methionine and gut SQOR deficiency synergize to cause intestinal injury

The interplay between dietary sulfur and metabolism on gut health was examined next (Fig. 2A). Mice were maintained on an isocaloric control (0.6% methionine) or 1.5% MRD. Serum thiosulfate was significantly elevated in IEC SQOR deficient mice on MRD (Fig. 2B). Genotype-associated differences in colon histology were not observed on the control diet (Fig. 2C, upper). However, a 10-day 1.5% MRD dietary intervention led to morphological changes in colon, which were more pronounced in SQOR-deficient mice (Fig. 2C, lower). Specifically, signs of crypt degeneration and decreased goblet cell density, and a marked thinning of the protective mucus layer were seen in SQOR deficient mice (Figs. 2D,E). MRD decreased levels of ATOH-1, a transcriptional regulator that commits intestinal stem cells to the secretory lineage (Figs. 2F,G), and MT-CO1 in SQOR-deficient mice (Fig. 2H,I). Interestingly, MRD attenuated the reductive shift in colon CoQ (Fig. 2J, Supplementary Figs. 2A,B), which was seen in SQOR-deficient mice on the control diet (Fig.1D). With the exception of mild isolated monocytopenia in SQOR-deficient mice, a complete blood count analysis did not reveal other MRD-induced differences between the genotypes (Supplementary Figs. 2C-I). Hemoglobin and hematocrit levels were also similar, indicating that the combination of gut sulfide oxidation deficiency and MRD did not lead to hematological changes (Supplementary Fig. 2J,K).

**Figure 2.**
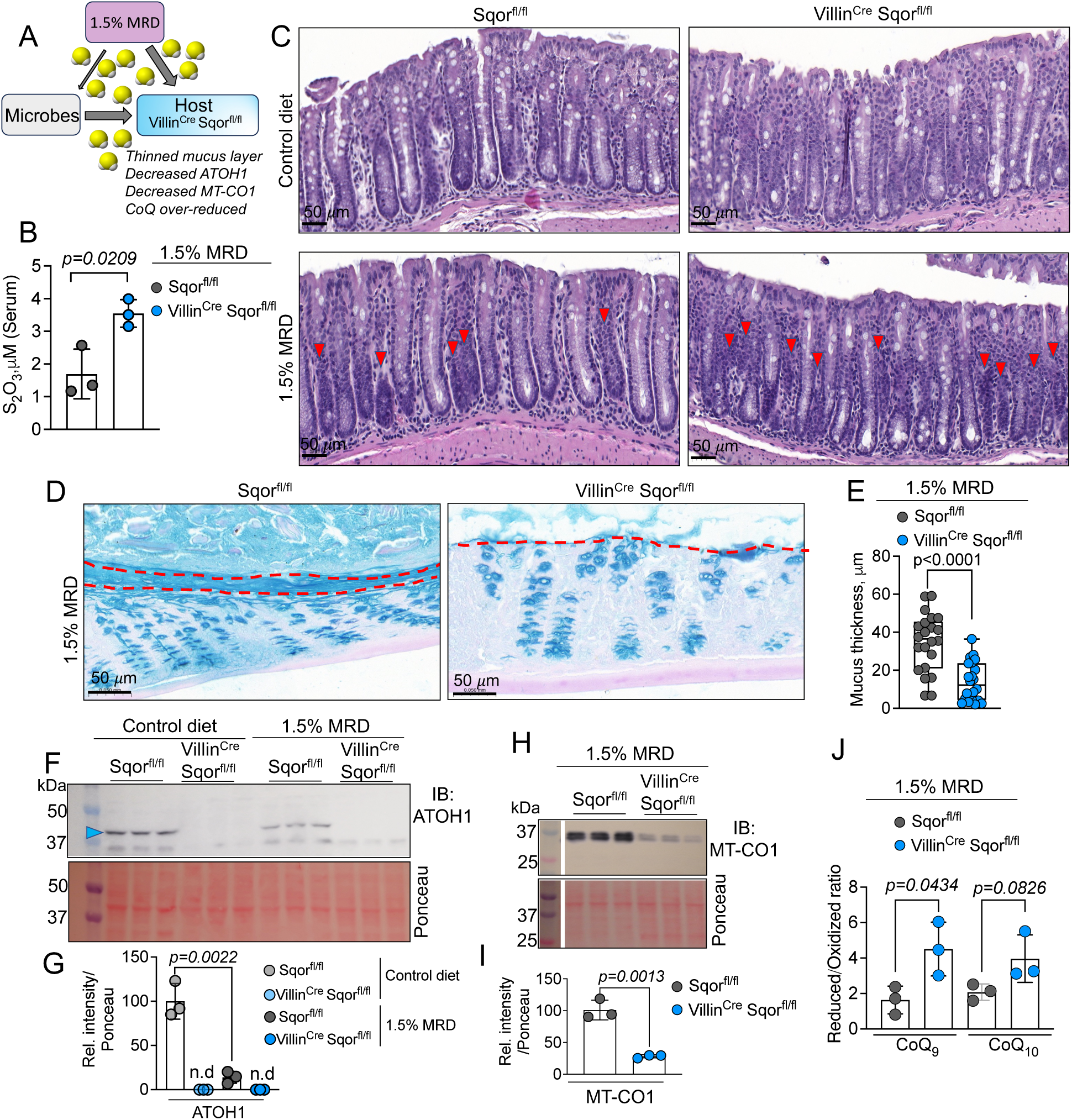
Genotype-diet modulation of colon architecture and redox. **(A)** Sulfide synthesis, impaired host sulfide oxidation, and MRD are predicted to synergize to induce intestinal injury in Villin^Cre^ Sqor^fl/fl^ mice. **(B)** Serum thiosulfate levels in the indicated groups (n=3 mice in each group). **(C)** Representative hematoxylin and eosin stained colon cross sections from Sqor^fl/fl^ and Villin^Cre^ Sqor^fl/fl^ mice on control versus 1.5% MRD (n=3). Red arrowheads point to diet-induced changes in crypt architecture**. (D-E)** Representative Alcian blue-PAS staining (D) and quantitative analysis (E) of mucin thickness in mice on a 1.5% MRD. (n=2 independent mice, 22 regions were used for quantitation of thickness). **(F-G)** Western blot (F) and quantitative analysis (G) of MT-CO1. (**H-I**) Western blot (H) and quantitative analysis (I) of ATOH1 in Sqor^fl/fl^ and Villin^Cre^ Sqor^fl/fl^ mice. (n=3 in F-I, nd is not determined). (**J**) Effect of 1.5% MRD on colon CoQ_9_ and CoQ_10_ redox ratio in the indicated genotypes (n=3).

### High dietary methionine and gut SQOR deficiency synergize to induce brain pathology

Patients with inherited SQOR deficiency present with neurologic Leigh disease-like lesions as seen by brain magnetic resonance imaging (MRI) (24). Given the significant contribution of gut H_2_S oxidation to systemic sulfide homeostasis (33), we asked if 10-11 week-old mice on control or 1.5% MRD exhibit brain pathology and/or behavioral changes after ∼6-8 weeks (Supplementary Fig. 3A). Diet-induced differences between the genotypes were not observed in weight, locomotor behavior, or grip strength (Supplementary Figs. 3B-D). T2-weighted MRI revealed areas of subtle hyperintensity and hypointensity (Supplementary Fig. 4A) which were similar to, albeit significantly less pronounced than changes seen in the whole-body SQOR mis-localization mouse model. Small Prussian blue positive foci suggested microbleeds (Supplementary Fig 4B). Unlike colon (Fig. 1B,C), brain MT-CO1 expression and CoQ_9_ and CoQ_10_ redox ratios were unaffected in mice lacking IEC SQOR, maintained on a control diet (Supplementary Fig. 5A-E).

Next, we challenged younger (∼5 week-old) mice with 1.5% MRD and performed biochemical behavioral, and imaging studies 11-15 weeks later (Fig 3A). Brain MT-CO1 levels were significantly lower in SQOR KO versus control mice (Fig 3 B,C) and the CoQ_9_ redox ratio was reductively shifted, albeit modestly (Fig. 3D and Supplementary Fig. 5F,G). These data reveal long-range effects on brain bioenergetics from the interaction between MRD and the IEC SQOR genotype. Remarkably, T2-weighted brain MRI revealed either the complete or asymmetric loss of lateral ventricles only in SQOR-deficient mice on MRD but not on the control diet (Fig. 3E and Supplementary Fig. 6). The diet-induced decrease in lateral ventricular volume indicated loss of cerebrospinal fluid (CSF). Transthyretin, a marker for choroid plexus where CSF is formed, was not significantly different (Fig 3F,G), while aquaporin 1, a water channel protein that plays a key role in CSF secretion, was significantly lower (Fig. 3H,I). Collectively, these data provide evidence for the long-range effects of loss of gut sulfide oxidation capacity on brain, which are sensitive to dietary methionine (Fig. 3J).

**Figure 3.**
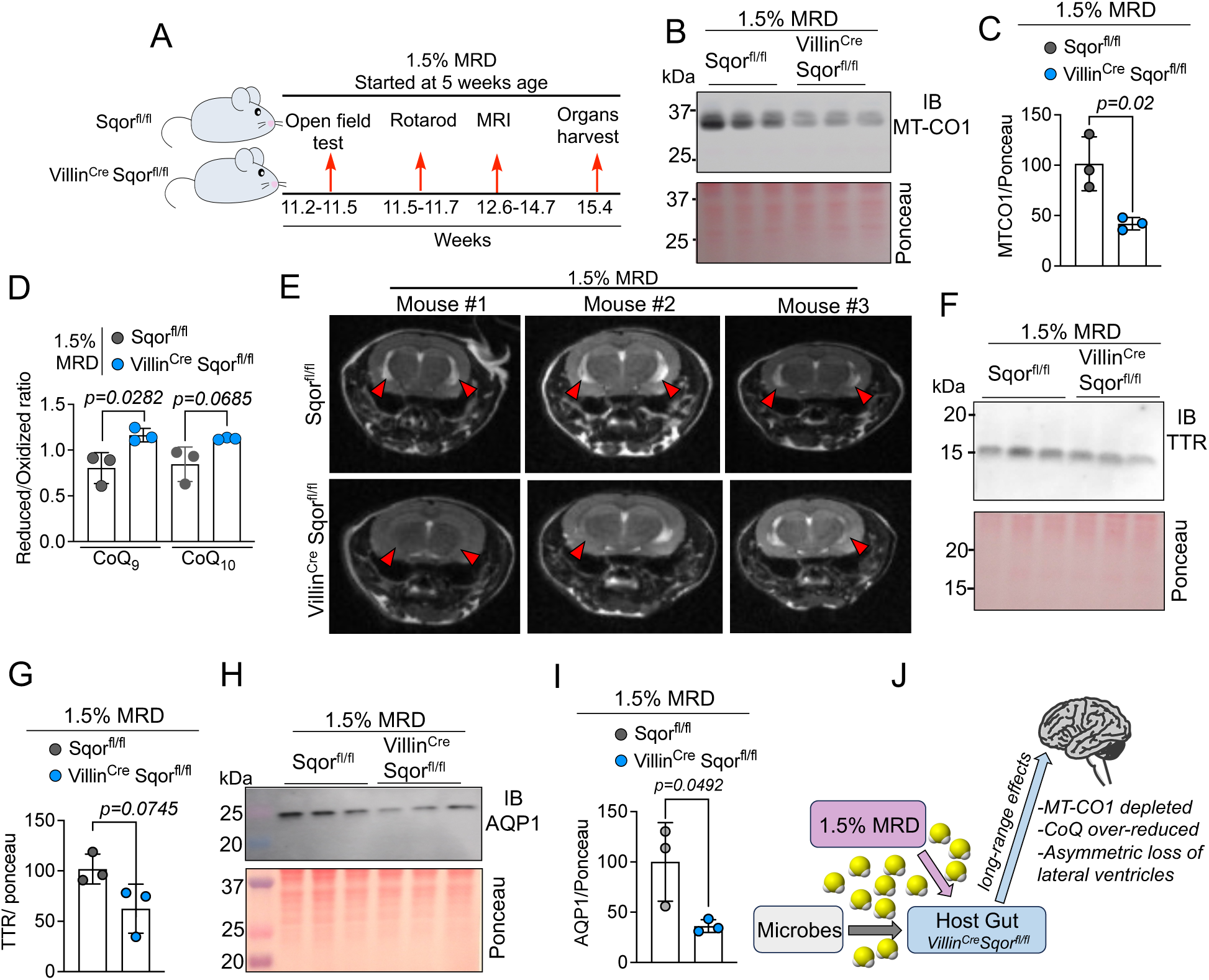
SQOR deficiency in gut epithelium perturbs brain CoQ redox and lateral ventricle structure. **(A)** Scheme showing experimental setup. (**B,C)** Western blot analysis (B) and quantitation (C) of MT-CO1 in murine brain on 1.5% MRD. n=3 independent mice for each conditions. **(D)** Brain CoQ redox status in mice maintained on 1.5% MRD (n=3 independent mice). **(E)** Representative axial T2-weighted MRI showing asymmetric or complete loss of lateral ventricles (red arrowheads) in Villin^Cre^ Sqor^fl/fl^ mice compared to controls, n=4 independent mice. **(F-I)** Western blot (F, H) and quantitation (G, I) of the choroid plexus markers, transthyretin (TTR), and aquaporin 1 (AQP1), n=3 independent mice. **(J)** Scheme showing long-range modulation of brain bioenergetics and function by diet and gut SQOR oxidation capacity.

### Alterations in microbiome composition and function in gut SQOR deficiency mice on MRD

Central to gut function and host homeostasis are the numerous microbes that make up the gut microbiome. To interrogate how diet interacts with IEC sulfide oxidation capacity, we profiled control and IEC SQOR-deficient mice on control diet versus MRD by 16S rRNA metagenomic sequencing of feces (Supplementary Fig. 7A). Alpha diversity, or the total number of taxa represented, was not significantly affected under any condition (Supplementary Fig. 7B). On the other hand, beta diversity, or the compositional makeup of the microbiome as measured by the Jaccard distance, was significantly different between SQOR-deficient versus control mice on MRD (Fig. 4A,B), but not between the other groups (Supplementary Fig. 7C-E). Among the significantly altered taxa, Lachnospiraceae, *Muribaculum* and *Chlamydia* were decreased while *Lactobacillus* and *Aldercreutzia* genera were present at a higher proportion in the microbiome of SQOR-deficient mice (Fig 4C,D).

**Figure 4.**
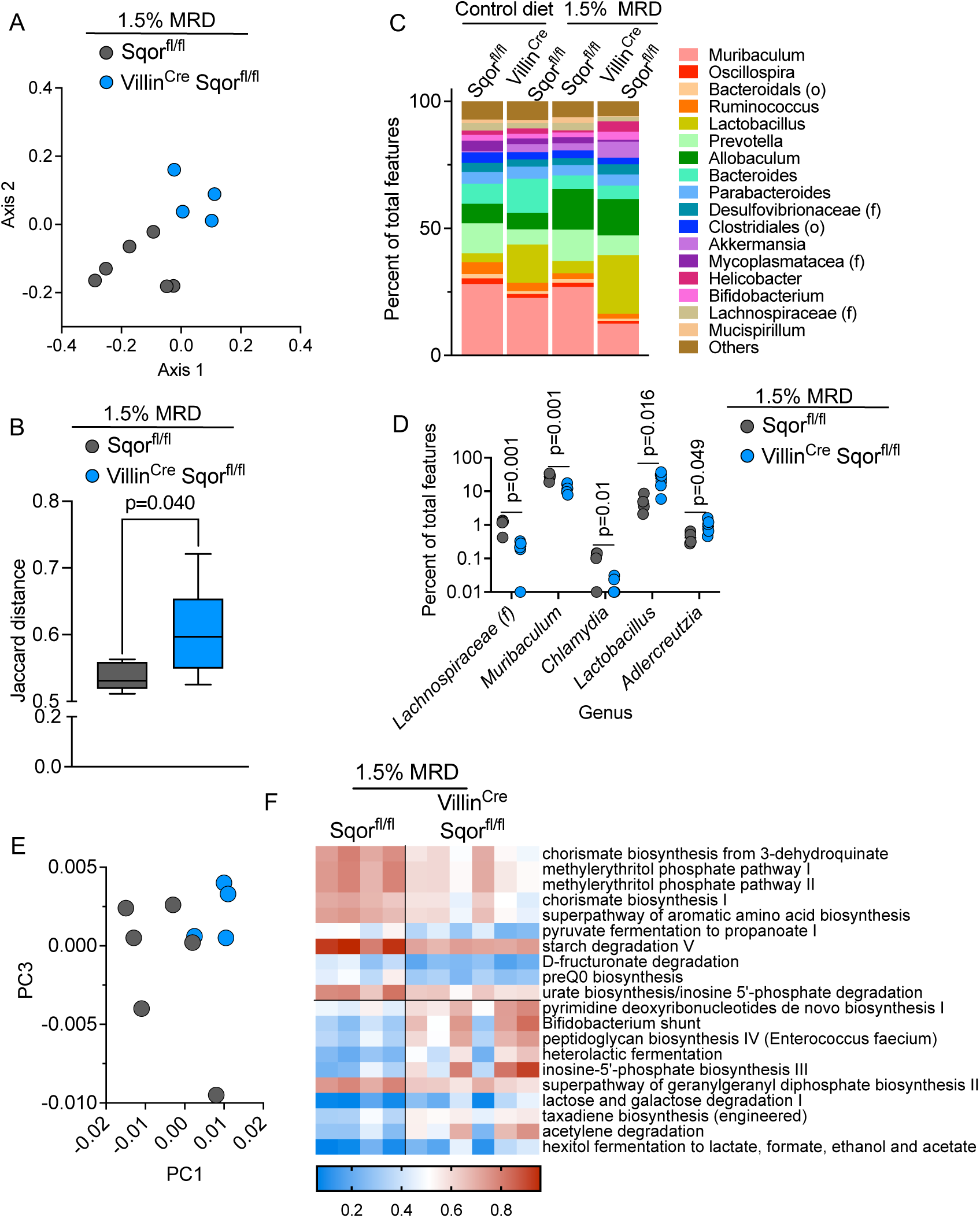
Microbiome composition and functional capacity are altered in IEC SQOR deficient mice on MRD. (**A**) Jaccard distance for the microbiome composition of control versus and IEC SQOR deficient mice, plotted on the first 2 principal component axes. (**B**) Jaccard distance within and between group pairwise comparison using PERMANOVA to calculate adjusted p value. (**C**) Composition of top 17 identified genera in each group. Taxa unable to be resolved beyond family (f) or order (o) are noted in the legend. (**D)** Significant differential genera between control and IEC SQOR deficient mice, as calculated by two-tailed t-test. (**E**) Principal component analysis of functional pathway representation in each microbiome, using functional metagenome prediction, and plotted on principal component axes PC1 and PC3. (**F**) Heatmap of top 10 increased and decreased pathways based on functional metagenome prediction.

Since the microbiome supports numerous host functions, including metabolite production that can impact brain physiology (35), we assessed how the observed compositional changes in the microbiome might impact its functional capacity. To this end, metagenomic functional prediction was employed on the 16S data and, as with the compositional differences, the SQOR-deficient group clustered away from control group on MRD (Fig. 4E). Among the most significant functions lost in SQOR-deficient mice were chorismate and aromatic amino acid biosynthesis, methylerithrytol phosphate metabolism, and pyruvate fermentation to propionate (Fig 4F, Table S1). On the other hand, SQOR-deficient mice had more functional representation of pyrimidine biosynthesis, fructose-6-phosphate fermentation and peptidoglycan biosynthesis (Fig 4F, Table S1). Together, these data indicate significant alterations in the microbiome in SQOR-deficient versus control mice on MRD, which could impact metabolite availability, and thereby, host homeostasis.

### Metabolomic signature of high methionine diet combined with gut SQOR deficiency

Serum metabolites were profiled for potential molecular clues underlying the long-range effects of 1.5% MRD on brain that are dependent on IEC SQOR deficiency (Fig. 5A,B, Table S2). Several interesting changes were seen in SQOR-deficient mice, including a decrease in tyrosine, which is a precursor of the catecholamines, epinephrine, norepinephrine and dopamine. Creatine, which is involved in high energy creatine phosphate metabolism particularly in skeletal muscle, was also decreased in SQOR-deficient mice together with the amino acids, aspartate and glutamate. In contrast, 5-hydroxyindole acetic acid, a catabolite of the hormone and neurotransmitter, serotonin, which is secreted primarily by enterochromaffin cells in gut, was increased. Additionally, the ketone bodies, acetoacetate and β-hydroxybutyrate were increased in SQOR-deficient mice. Since liver is a primary organ for methionine metabolism and creatine and ketone body synthesis, MT-CO1 levels were examined in this organ as a marker of H_2_S-induced ETC dysfunction; they were lower in SQOR-deficient mice on 1.5% MRD (Fig. 5C,D). Liver thiosulfate, a proxy marker for H_2_S exposure, was significantly higher is SQOR deficient mice (Fig. 5E). Impaired oxidative phosphorylation in liver is expected to favor acetyl-CoA flux towards ketone body synthesis over the TCA cycle, whereas ketolysis occurs in peripheral tissues but primarily in brain, heart and skeletal muscle (Fig. 5F).

**Figure 5.**
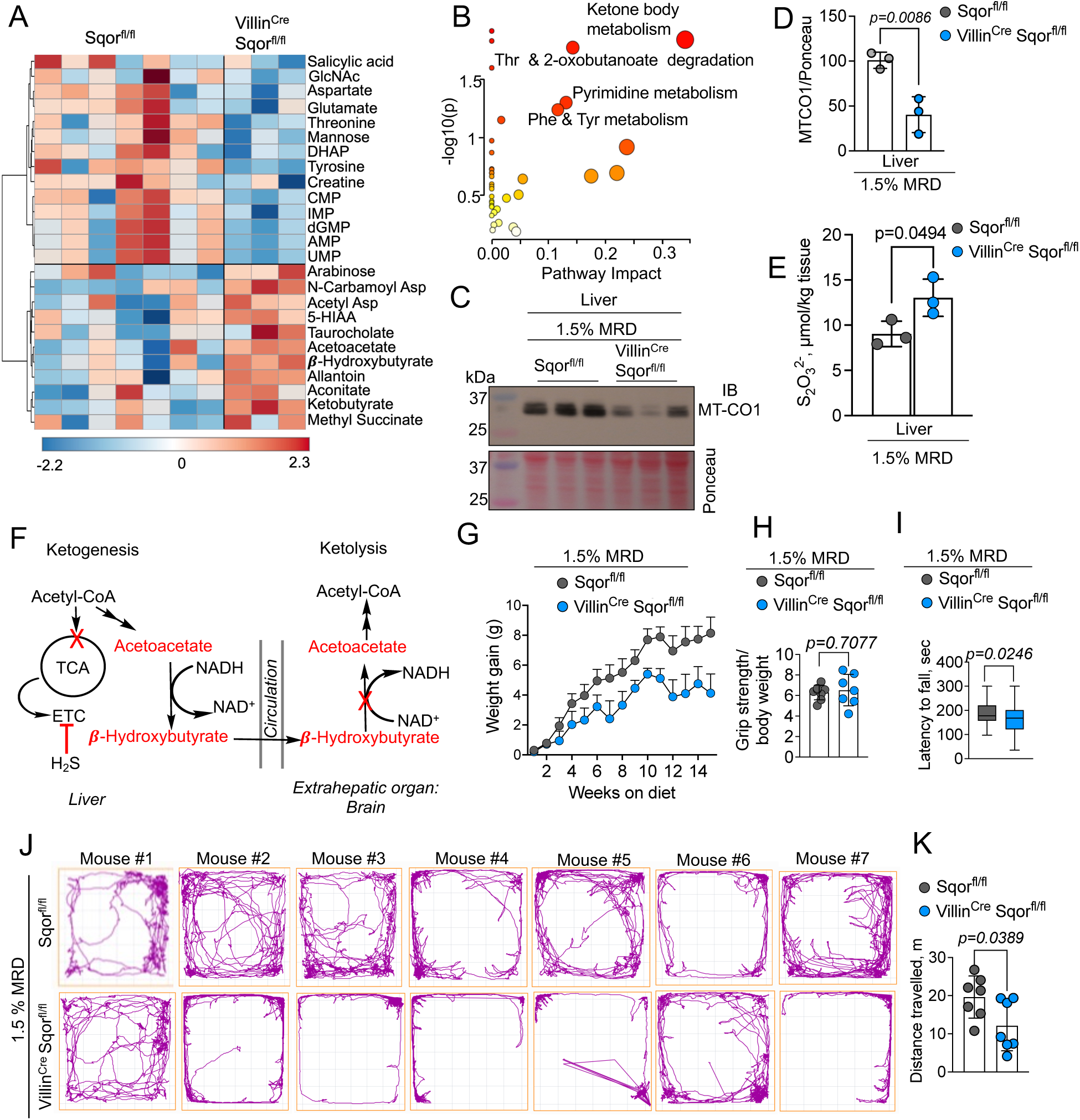
Impaired gut sulfide oxidation promotes ketosis and impairs motor coordination. **(A)** Heat map showing significant changes in serum metabolites in Sqor^fl/fl^ and Villin^Cre^ Sqor^fl/fl^ mice on 1.5% MRD. **(B)** Pathway analysis by Metaboanalyst showing that ketone body metabolism is most impacted. **(C,D**) Western blot showing lower MT-CO1 levels (C) and quantitation (D), n=3 independent mice. **(E)** Liver thiosulfate levels in Sqor^fl/fl^ and Villin^Cre^ Sqor^fl/fl^ mice. (**F**) Scheme showing sulfide induced ketogenesis in liver leading to increased ketone bodies in circulation. Ketolysis occurs in extrahepatic tissue like brain. **(G)** Weight change in Sqor^fl/fl^ and Villin^Cre^ Sqor^fl/fl^ mice on 1.5% MRD. **(H, I)** Changes in (H) grip strength and motor coordination assessed by the rotarod test (I). **(J, K)** Differences in movement trajectories (J) and distance traveled (K) as monitored in an open field test, n=7 independent mice.

### High dietary methionine and gut SQOR deficiency elicit motor defects

When introduced to MRD at 5 weeks, mice lacking IEC SQOR exhibited lower weight gain over 15 weeks compared to floxed controls (8.1 ± 3.0 versus 4.1 ± 3.4 g, Fig. 5G). While a difference in grip strength was not observed between the genotypes (Fig. 5H), the rotarod test, which measures resistance to fatigue while gripping a rotating rod, revealed significant differences (Fig. 5I). In the open field test to measure exploratory locomotor activity, SQOR-deficient mice traveled significantly shorter distances and expressed lower curiosity for exploration than control mice (Figs. 5J,K). A profound difference in exploratory locomotor activity was also apparent by simply observing the mice (Movie S1).

## DISCUSSION

Our study reveals that the interaction between diet and sulfide metabolism at the host-gut interface can have long-range effects on liver, brain and behavior. Protection from high luminal exposure to H_2_S is afforded by high expression of SQOR in colon (20), and depends on CoQ availability and recycling of its reduced form by complex II and/or III (33). SQOR loss in IECs has a profound effect on the CoQH_2_:CoQ ratio, which shifts from ∼2 to ∼10, and, remarkably, is corrected by broad spectrum antibiotics treatment (Fig. 1F,G). The CoQ redox shift is consistent with complex IV inhibition due to H_2_S accumulation, and is correlated with loss of the MT-CO1 subunit in cytochrome c oxidase (Fig. 1B,C). The drop in MT-CO1 mirrors the response of colon adenocarcinoma cells exposed to sulfide, which is accompanied by inhibition of respiration (17). MT-CO1 depletion has also been observed in a murine model of ethylmalonic encephalopathy, which is associated with sulfide accumulation (32). As with the CoQ redox status, antibiotics restore MT-CO1 levels (Fig. 1E,F). The correction of CoQ redox and MT-CO1 levels help explain the effect of metranidozole on restoring complex IV activity in whole-body SQOR-deficient mice (26). The phenotypic reversibility seen in our study provides strong evidence for the impact of microbial-derived sulfide on gut bioenergetics, and predicts a vulnerability to intestinal injury when sulfide oxidation capacity is compromised.

Since CoQ serves as an electron acceptor for multiple enzymes that intersect with the ETC, a reductive shift would also adversely impact other users, including the electron transfer flavoprotein quinone oxidoreductase. The latter serves as a conduit for electrons in the β-oxidation of butyrate, the preferred fuel for differentiated colonocytes (36). Disrupted butyrate utilization would disturb the butyrate gradient in crypts and inhibit stem cell proliferation (37). A reductive shift is expected to ripple out from CoQH_2_ to the NADH and FADH_2_ pools, which transfer electrons to CoQ at complexes I and II, respectively. On the other hand, CoQH_2_ accumulation is predicted to promote its antioxidant function, protecting against lipid peroxidation (38-40). The clinical manifestations of hereditary CoQ deficiency include nephropathies, myopathies and ataxias (41). The potential effects of a secondary electron acceptor insufficiency due to an over-reduced CoQ pool are however, not known.

Despite changes in mitochondrial ETC markers (i.e., CoQ redox and MT-CO1 levels), SQOR deficiency in IECs did not visibly impact crypt architecture or goblet cell density (Fig. 2C). However, when the mice were challenged with 2.5-fold higher methionine relative to an isocaloric diet, the genotypic liability was revealed within 10 days. Degraded crypts, decreased goblet cell number, and a significant thinning of the mucus layer were visible (Fig. 2C-E). The mucus layer is a network of mucins secreted by goblet cells, and serves as a lubricant and a defensive barrier, limiting host exposure to luminal content. Mucin polymers are stabilized by disulfide bonds and form a hydrated gel-like inner layer that is sterile, and a loosely interconnected outer layer that is penetrated by microbes (42). Thinning and decreased mucus coverage are associated with various intestinal pathologies including cancer, inflammatory bowel disease and enteric infections (42). While the intrinsic reactivity of sulfide with disulfides is low (0.6 M^-1^ s^-1^, pH 7.4, 20 °C) (43), sulfide accumulation due to changes in diet and/or compromised sulfide oxidation capacity, or microbial composition, could potentially enhance disulfide cleavage, a postulated mechanism for mucus barrier degradation in inflammatory bowel disease (44). Thinning of the mucus layer could result from fewer goblet cells, impaired secretion, and/or structural destabilization of mucin. The significantly lower expression of ATOH1, a master transcriptional regulator in intestinal stem cells of the secretory lineage, correlates with the decrease in goblet cell numbers (Fig. 2B,E). Elevated serum thiosulfate, provides strong evidence for increased H_2_S exposure, which is further supported by lower MT-CO1 and a reductively shifted CoQ_9_ pool (Fig. 2B, H-J).

The microbiome is responsible for much of the metabolic activity in gut, which can impact host metabolism systemically (45). Microbiome analysis revealed that IEC SQOR deficiency potentiated the effects of MRD, leading to compositional differentiation from the control group. Changes in important taxa such as notable butyrate producers in the Lachnospiraceae family (46) and immunomodulatory *Lactobacillus* species (47) were observed (Fig. 4D). Functional metagenomic analysis predicted downregulation of chorismate biosynthesis and aromatic amino acid metabolism, which are associated with neurotransmitter as well as tyrosine synthesis (48); the latter was decreased in serum of IEC SQOR-deficient mice. The methylerythritol phosphate pathway genes play key roles in oxidative stress response (49) and were also predicted to be downregulated. We posit that the microbiome changes influence MRD-induced systemic effects in IEC SQOR-deficient mice.

The typical neuroimaging finding in Leigh disease is symmetrical hyperintense loci in brain stem or basal ganglia, although some patients display other pathologies, including cerebellar lesions, delayed myelination or cerebral atrophy (25). Subtle indications of symmetric hyperintense regions as seen in Leigh disease, were observed in mice lacking IEC SQOR and maintained on MRD for 6-7 weeks (Supplementary Fig. 4A). While the genotype *per se* did not affect brain CoQ redox status (Supplementary Fig. 5C), it synergized with MRD to induce a reductive shift (Fig. 3D). When the mice were maintained on MRD for ∼13-15 weeks, asymmetric or complete loss of lateral ventricles was seen by MRI (Fig. 3E), which, to our knowledge, has not been reported for Leigh disease.

In humans, asymmetric lateral ventricles without a clear pathological cause (e.g. intracranial bleeding, infarction, trauma or space-occupying lesions) is seen in 5-12% of healthy individuals (50, 51). AQP1 is the principal water channel found in the apical side of the choroid plexus epithelium, and its loss is correlated with diminished CSF production and ventricle size (52). IEC SQOR-deficient mice on MRD showed ∼3-fold lower AQP1 protein levels than littermate controls (Fig. 3H,I). H_2_S has been previously reported to downregulate AQP5 expression in lung (53), while persulfidation of AQP8 at a specific cysteine residue reportedly inhibits H_2_O_2_ transport in HeLa cells (54).

Ketone body metabolism was perturbed by IEC SQOR deficiency on MRD and serum acetoacetate and β-hydroxybutyrate were found to be elevated (Fig. 5A,B). SQOR deficiency patients presented with elevated ketone levels in urine (24). Ketogenesis is promoted under conditions of glucose or insulin insufficiency and/or decreased TCA cycle flux. Decreased hepatic MT-CO1 in SQOR-deficient mice (Fig. 5C,D) is indicative of restricted ETC capacity in liver, while thiosulfate levels are consistent with higher H_2_S exposure (Fig. 5E). Ketone bodies serve as a primary fuel for brain during fasting or starvation, but can also be diverted to lipid and sterol synthesis or excreted in urine. Additionally, non-canonical signaling functions have been ascribed to ketone bodies, e.g., β-hydroxybutyrate inhibition of class I HDAC increases histone acetylation (55). Similar biochemical responses to sulfide exposure seen in cell culture, i.e., decreased MT-CO1 levels (17) and increased thiosulfate (56), supporting our conclusion that the changes observed in liver and brain resulted from increased circulating sulfide.

A limitation of our study is the single time point for brain MRI analysis. While brain MRI on mice maintained on MRD for 6-7 weeks revealed very subtle Leigh disease-like pathology, it would be informative to assess whether the lesions progress with age in the IEC SQOR deficiency model as seen in the mis-localized SQOR mouse model and in patients (24, 26). Second, the increase in dietary methionine has other metabolic consequences, including an increase in homocysteine, which is itself, a substrate for H_2_S biogenesis (6) but also has other effects, which were not studied. Finally, the impact of diet and SQOR deficiency on gut microbial composition or genotype-diet interactions in the presence of antibiotics were not addressed. The current study lays the groundwork for these future interrogations.

In summary, we demonstrate that loss of sulfide oxidation capacity in gut epithelial, increases sulfide exposure systemically, and elicits both local as well as long-range effects. Locally, the combination of a 2.5-fold increase in dietary methionine and SQOR deficiency degraded crypt architecture, decreased goblet cell density, and compromised mucus layer thickness, which could increase the propensity for inflammation and intestinal diseases (57). The genotype-diet interaction also decreased liver and brain ETC capacity, and increased ketogenesis, signaling perturbations in energy utilization. Brain MRI in mice IEC SQOR KO mice on MRD, revealed loss of ventricular volume, which was associated with lower AQP1, important for CSF secretion. Our data provide intriguing evidence for systemic perturbations in energy metabolism and behavior that originate from loss of sulfide oxidation capacity in gut in combination with elevated dietary methionine. Our data suggest that restricted methionine intake could be beneficial for treating diseases resulting from compromised sulfide oxidation capacity, e.g., due to loss of SQOR or ETHE1. Focal loss of SQOR in IECs represents a potentially useful model for analysis of the three-way gut microbe-host-diet interaction in health and disease.

## ACKNOWLEDGEMENTS

This work was supported in part by the grants from the NIH (National Institutes of Health R35GM130183 to RB, R01CA248160 to CAL and R01CA148828 and R01CA245546 to YS), the Division of Intramural Research, National Institute of Allergy and Infectious Diseases, NIH (to SKH), the American Heart Association (826245 to RK), the American Physiology Society and Crohn’s and Colitis Foundation (1003279 to RS). We thank Dr. Nupur Das (University of Michigan) for guidance with sample collection for microbiome analysis.

## DATA AND MATERIALS AVAILABILITY

All data are available in the manuscript or supplementary materials.

## COMPETING INTEREST STATEMENT

C.A.L. has received consulting fees from Astellas Pharmaceuticals, Odyssey Therapeutics, and T-Knife Therapeutics, and is an inventor on patents pertaining to Kras regulated metabolic pathways, redox control pathways in pancreatic cancer, and targeting the GOT1-pathway as a therapeutic approach (US Patent No: 2015126580-A1, 05/07/2015; US Patent No: 20190136238, 05/09/2019; International Patent No: WO2013177426-A2, 04/23/2015).

## AUTHOR CONTRIBUTIONS

RK, YS and RB conceptualized the study and RK performed and analyzed the majority of the experiments and was assisted by: DJS-open field test, VIB-microbiome analysis, MLS-Alcian blue PAS staining, PS, HW and CAL-metabolomics data collection and analysis, RS-RT-qPCR, VV-liver thiosulfate determination, FI-technical advice on brain MRI data acquisition, SKH-resource support. RK and RB drafted the manuscript and all authors edited and approved the final version.

## Methods

### Materials

Antibodies were purchase from the indicated vendors. MT-CO1 (1D6E1A8) and MT-CO2 (ab110258) were from Abcam and ATOH1 (21215-1-AP), transthyretin (11891-1-AP), aquaporin 1 (20333-1-AP), IBA 1 (10904-1-AP), GFAP (60190-1-Ig), CD31 (28083-1-AP), SQOR (17256-1-AP) were from Proteintech. Secondary anti-rabbit horseradish peroxidase-linked IgG antibody (NA944V), and secondary anti-mouse horseradish peroxidase-linked IgG antibody (NA931V) from GE healthcare.

Reagents were purchased from the indicated vendors. Hematoxylin (Newcomer Supply), eosin, Alcian blue, periodic acid (Alfa Aesar), Schiff’s reagent (Electron Microscopy Sciences), Carnoy’s fixative (Spectrum), xylene, paraffin, ethanol, buffered formalin (Fisher Scientific, SF98-4), CoQ_9_, CoQ_10_, methanol, isopropanol, hexane, *n*-propanol, *p*-benzoquinone, sodium borohydride, protease inhibitor cocktail for mammalian tissue extract (P8340), RIPA lysis buffer (R0278), SuperScript III One-Step RT-PCR System with Platinum Taq DNA Polymerase (12574026, Invitrogen) and SYBR Green PCR master mix (4309155, Applied Biosystems), penicillin G, ampicillin, neomycin, gentamycin, vancomycin, metronidazole, PVDF membrane, Kwik Quant substrate, Biorad substrate, phosphate buffered saline, sodium thiosulfate (217263), monobromobimane (M20381, Invitrogen), Tris-base, meta-phosphoric acid, methionine rich diet (A10021Bi and A21111101i, Research Diet, INC.), and Ponceau (Sigma, P3504).

### Animal studies

Mice were maintained under standard housing conditions (22 ± 2 °C, humidity ranging from 30% to 70% and 12-h light/dark cycle). Mice were provided with ad libitum access to food and water and all experiments were performed with approval from the Institutional Animal Care and Use Committee at the University of Michigan.

### Villin^Cre^ Sqor^fl/fl^ knockout mice

Sqor^loxP/+^mice were generated by Cyagen Biosciences as described previously (15). An intestinal epithelial disruption of Sqor was generated by crossing Sqor^loxP/loxP^ mice to a Villin-Cre line (58).

### Antibiotics treatment

Antibiotics treatment was performed as described previously (59). A Briefly, mice were given drinking water containing an antibiotic cocktail (ampicillin 1 g/L, neomycin 1 g/L, gentamycin 500 mg/L; penicillin 100 U/L). In addition, vancomycin (1 mg/mL) and metronidazole (0.5 mg/mL) were introduced via oral gavage on alternate days for 10 days.

### CoQ redox measurement

Harvested organs were dropped in liquid nitrogen, frozen within a couple of minutes and transferred to cryotubes for storage at −80°C until use. Frozen mouse tissues (colon and brain) were homogenized in 4-6 volumes of n-propanol with a glass homogenizer. The lysates were centrifuged at 12,000 x g for 5 min at 10°C and the supernatant was collected carefully without disturbing the pellet, and stored at −80 °C until use. Samples were analyzed within 2-weeks of supernatant collection. HPLC analysis of CoQ was performed on a Microsorb-MV 100-5 C18 column (150 × 4.6 mm, 5 m bead size, Agilent) as described previously (34). Briefly, samples were eluted isocratically with solvent, comprising isopropanol (15 mL), methanol (845 mL), and hexane (140 mL) in a total volume of 1L, with a flow rate of 0.8 mL/min at room temperature. Peaks were detected by UV absorbance at 275 nm. Oxidized and reduced CoQ exhibit absorbance maxima at 278 nm (ε = 14,500 M^−1^ cm^−1^) and 287 nm (ε = 3340 M^−1^ cm^−1^) respectively, and calibration curves were generated with oxidized CoQ_9_ and CoQ_10_ samples of known concentration prepared in n-propanol. Each sample was run twice, i.e., before and after oxidation of reduced CoQ with *p*-benzoquinone. Samples were reduced by mixing the supernatant (86 µL) with *p*-benzoquinone (86 µL, 4 mg/mL) for 5-8 mins at 10 °C in a autosampler before being injected onto the HPLC column. Sample analysis was done as described previously (34).

### Western blotting

Mouse tissue was homogenized in RIPA buffer using a bead homogenizer (TissueLyzer III, Qiagen) and the supernatant was collected after centrifugation at 12,000 x *g* for 5 min at 4 °C and stored at -80°C until use. The Bradford assay was used to quantify protein in the supernatant, which was then diluted with 4X Laemmli sample buffer (62.5 mM Tris-HCl, pH 6.8, 10% glycerol, 1% LDS, 0.005% bromophenol blue) to obtain a final concentration of 2 µg of protein per µL. Samples were heated at 95 °C for 5 min and used immediately for western blotting or stored at −80 °C for upto 4 weeks. Proteins were separated on 12% SDS polyacrylamide gels and transferred to an activated PVDF membrane, using a Bio-Rad trans-blot turbo transfer system. Membrane was blocked with 5% skimmed milk and incubated overnight at 4 °C with primary antibodies with the following dilutions: MT-CO1 (1:1000), SQOR (1:2000), ATOH1 (1:1000), CD31 (1:500), AQP1 (1:5000), GFAP (1:1000), and IBA1 (1:1000). The secondary antibody, horseradish peroxidase-linked anti-rabbit IgG and anti-mouse IgG were used at a 1:10,000 dilution. Blots were developed with KwikQuant Digital-ECL substrate (KwikQuant), and images were collected with a KwikQuant Imager. Equal loading was verified by Ponceau S staining of membranes post chemiluminescent detection of signals.

### Quantitative RT-PCR

Total RNA was extracted from colon tissue using a Trizol. RNA (500 ng) was reverse transcribed to cDNA using a SuperScript III First-Strand Synthesis System (Invitrogen). The following primer pairs were used for amplification: 5′-GAGTTGGAGCAGAGAATGTGGC-3′ and 5′-CACACTCAGTGTGGAACGGACT-3′ (SQOR) and 5′-GGCTGTATTCCCCTCCACG-3′ and 5′-CCAGTTGGTAACAATGCCATGT-3′ (ACTB, cytoplasmic actine). The cDNA, gene-specific primers and SYBR Green master mix (Applied BioSystems) were combined and run in a QuantStudio 5 Real-Time PCR System (Applied BioSystems). Relative gene expression was estimated using the change in cycling threshold (ΔΔCt) method with ACTB, serving as a housekeeping gene.

### Histological analysis

Following necropsy, intestines were harvested, rinsed, and rolled into Swiss rolls followed by incubation in 10% buffered formalin phosphate (Fisher Chemical) overnight at room temperature. Samples were then incubated in 70% ethanol for 24-72 h and then embedded in paraffin. For mucus barrier analysis, the intestine was excised, and horizontal cuts were made above and below a fecal pellet. Samples were incubated at room temperature overnight in Carnoy’s Fixative (Spectrum) to preserve mucus integrity. Next, samples were transferred to 70% ethanol and stored at 4°C for 24-72 h until paraffin embedding and cross-sectioning.

*Hematoxylin and eosin staining-*Tissues were fixed with PBS-buffered formalin for 24 h, followed by embedding in paraffin. Sections of 5 μM thickness were stained with H&E and mounted with Permount Mounting Medium.

*Alcian blue periodic acid Schiff (PAS) staining-*Following paraffin wax embedding and sectioning, histological samples were deparaffinized with 99% xylene and rehydrated with decreasing dilutions of ethanol (100% and 95%) and water. Samples were incubated for 3 min in 3% acetic acid and then in Alcian Blue (Newcomer Supply) for 30 min at room temperature. Samples were oxidized for 10 min in 0.5% periodic acid (Alfa Aesar), followed by a 20-min incubation in Schiff’s reagent (Electron Microscopy Sciences). Samples were dehydrated using ethanol (95% followed by 100%) and 99% xylene and coverslips were mounted on samples using Permount (Fisher Chemical).

*Prussian Blue Staining for Iron-*The Abcam iron stain kit (ab150674) was used to detect iron in mouse brain sections. Whole brain was embedded in O.C.T compound (Fisher Healthcare), sectioned in 10 μm slices using a cryostat, and stored at -80°C. Staining was performed within a month of sectioning. On the day of staining, brain sections were brought to room temperature and incubated in 1x PBS, pH 7.4, for 4 min. Then, the PBS was carefully removed with a pipette, and the sections were incubated for 3 min in a 1:1 solution of potassium ferrocyanide and hydrochloric acid provided in the kit. The slides were subsequently washed with distilled water and incubated in a nuclear fast red solution for 5 min. After another wash with distilled water to remove excess nuclear fast red, the sections were dehydrated using 95% ethanol and mounted with an acrylic mounting medium (Newcomer supply) and left to dry overnight at room temperature. Imaging was performed using a slide scanner.

### Brain MRI

Mice were anesthetized with 2% isoflurane/air mixture throughout MRI examination. Mice lay prone, head first in a horizontal bore 7.0T Bruker MR scanner with the body temperature maintained at 37 °C, using a circulating heated water bath. A quadrature volume radiofrequency coil was used to scan the head region of the mice. Axial images were acquired using a T2-weighted rapid acquisition with relaxation enhancement (T2-RARE) sequence with the following parameters: repetition time (TR)/effective echo time (TE), 4000/60 msec; echo spacing, 15 msec; number of echoes, 8; field of view (FOV), 20 x 20 mm; matrix, 256 x 128; slice thickness, 0.5 mm; number of slices, 25; and number of scans, 1 (scan time 8.5 min).

### Rotarod Test

Mice were placed on a stationary rod (1.25 inch diameter) of rotarod from IITC Rotarod Series 8. If mice fell off before the rotation began, they were placed back on the rod. If mice fell off repeatedly, indicating a problem, the animal was not used in the experiment. With mice successfully perched on the rod, the rotation was initiated. The rod was programmed to begin rotating at 4 rpm and increase to 45 rpm in 300 sec. When an animal fell off the rod, it landed on a tray below triggering the program to record the final elapsed time (s), distance traveled (m) and speed obtained (rpm). This procedure was repeated after the animal had rested for a few minutes.

### Grip Strength

With the meter (Columbus Instruments Grip Strength Meter model 0167-005L) set to measure Kg Peak Tension and reset to zero, mice were carefully lifted by their tails so that they grasped the horizontal bar firmly with both paws. Then, in a firm, but gentle, and constant motion, the animal was pulled pull away until its grasp is broken. The displayed peak tension value was recorded. Each animal was tested in three trials, and the peak tension value was averaged and normalized to body weight.

### Open field test

The open-field test was originally developed in 1934 to test emotionality in rodents (60). We used the open-field assay to test locomotor and anxiety-like behavior, using a modified version of a validated and published protocol (60). A total of 14 animals were tested using the open-field assay; half were controls (n=7) and the other half were (n=7) SQOR knockouts. The Stoelting Company open-field apparatus (40L x 40W x 40H) is broadly used to test exploratory behavior in mice. The apparatus has a grey metal base that detaches from the walls for cleaning with 70% ethanol between trials to decrease confounding behavior with olfactory cues left by mice used in the previous measurement.

The walls of the apparatus were covered with opaque paper to reduce interference with visual cues. After placing the mouse in the apparatus and toggling the start button for recording, both observers left the room. The mice moved freely in an open field apparatus for 5 min. The distance moved was quantified using ANY-Maze tracking software, version 7.4. with a V500s 8MP doc camera connected to a computer. A camera was placed above the open field arena for recording. ANY-Maze was also used to generate track plots for qualitative data tracing animal movement in the open field arena.

### Serum metabolomics

Blood was harvested from submandibular vein in 1.5 mL Eppendorf tube and allowed to stand at room temperature for 30-45 mins then centrifuged at 3000 x g for 15 min at 4 °C and supernatant was stored at -80 °C. Within 2 days, frozen samples were thawed on ice and 50 µL of samples were mixed with 200 µL of 100% ice cold methanol (80% v/v methanol). Next, the samples were vortexed and incubated on dry ice for 10 min. Samples were centrifuged at 13,000 x g for 10 min at 4 °C and the equal volume of supernatant was transferred to 1.5 mL Eppendorf tube and submitted to metabolomics core facility at University of Michigan where data were acquired as described previously (61).

### Serum and tissue thiosulfate quantitation

Serum was prepared as described above in the serum metabolomics section and stored at -80°C. Serum (22.5 μL) was mixed with 1.25 μL of 1 M Tris base and 60 mM MBB and protein was precipitated using metaphosphoric acid (MPA, 50 μL, 16.8 mg ml^-1^). Frozen tissue samples were ground on ice with 4 volumes of 20 mM HEPES buffer, pH 7.4, and 40 μL of homogenate was placed in a separate tube containing 2.5 μL 1M Tris and frozen in dry ice and then mixed with 7.5 μL of 60 mM MBB. Tubes containing tissue homogenate with Tris and MBB were thawed, vortexed and incubated at room temperature for 10 min in the dark. Proteins were precipitated with MPA as described above. Derivatized and precipitated serum and tissue homogenate were centrifuged at 12,000 x *g* for 5 min at 4 °C. The supernatants were collected in the dark and stored at −20 °C until further use. Derivatized samples were analyzed within 2 days by HPLC. A C18 column (Agilent, Zorbax Eclipse XDB, 5 μm, 4.6 × 150 mm) was used to separate thiosulfate using an ammonium acetate/methanol buffer system as described previously (62).

### Microbiome analysis

For bacterial community analysis, mouse fecal samples were obtained by gently holding each mouse over a clean Petri dish within a laminar flow hood to encourage natural defecation. The samples were promptly placed in sterile tubes, quickly frozen on dry ice, and stored at -80°C for up to two week before analysis. Bacterial sequencing analysis was carried out at the University of Michigan Host Microbiome Core facility. The V4 region of the 16S rRNA gene was amplified from each sample utilizing a dual-indexing sequencing strategy. PCR amplification conditions were as follows: initial denaturation at 95°C for 2 min, followed by 30 cycles of 95°C for 20 sec, 55°C for 15 sec, and a final extension at 72°C for 10 min. Accuprime High Fidelity Taq (ThermoFisher, Grand Island, NY) was used for amplification. The PCR amplified samples were frozen at -20 °C until further use. Amplicons were visualized on an eGel 96 SYBR Safe 2% gel system (ThermoFisher) and normalized with the SequalPrep Normalization Plate Kit (ThermoFisher). Sequencing was performed on the Illumina MiSeq platform with the MiSeq Reagent Kit V2 (500 cycles), following the manufacturer’s protocol.

### Hematological analysis

Complete blood count analysis was conducted by the Unit for Laboratory Animal Medicine Pathology Core at the University of Michigan.

### Statistical Analysis

GraphPad Prism 10 was used for statistical analysis, and the p-values from two-sided Student’s t-tests are indicated in the figure panels. Analysis of 16S sequencing was performed using QIIME2 (63). Amplicon sequence variants were generated with DADA2 and reads were aligned to the Greengenes13_8 reference set for taxonomic classification. Functional metagenomic inference was accomplished using Picrust2 (64) on the QIIME2 output data. These data were analyzed with STAMP software (65).

