## Supplementary information for "Gut sulfide metabolism modulates behavior and brain bioenergetics"

**Table of Content**

**Supplementary Figure 1.** Validation of Sqor KO and CoQ quantitation
**Supplementary Figure 2.** Complete blood count and CoQ analysis
**Supplementary Figure 3.** Effect of MRD on Villin$^{Cre}$ Sqor$^{fl/fl}$ mice
**Supplementary Figure 4.** Effect of MRD on brain MRI in Sqor$^{fl/fl}$ and Villin$^{Cre}$ Sqor$^{fl/fl}$ mice
**Supplementary Figure 5.** Effect of MRD on brain bioenergetics in control versus Villin$^{Cre}$ Sqor$^{fl/fl}$ mice
**Supplementary Figure 6.** Comparison of T2-weighted brain MRIs
**Supplementary Figure 7**. Alpha diversity and beta diversity comparisons in control versus Villin$^{Cre}$ Sqor$^{fl/fl}$ mice on control or 1.5% MRD
**Table S1**.Sheet 1:Significant microbiome pathways (sheet 1), microbiome composition (sheet 2) and raw data for microbial pathway analysis (sheet 3)
**Table S2.** Serum metabolomics data
**Movie S1.** Activity of Sqor$^{fl/fl}$ versus Villin$^{Cre}$ Sqor$^{fl/fl}$ mice on MRD

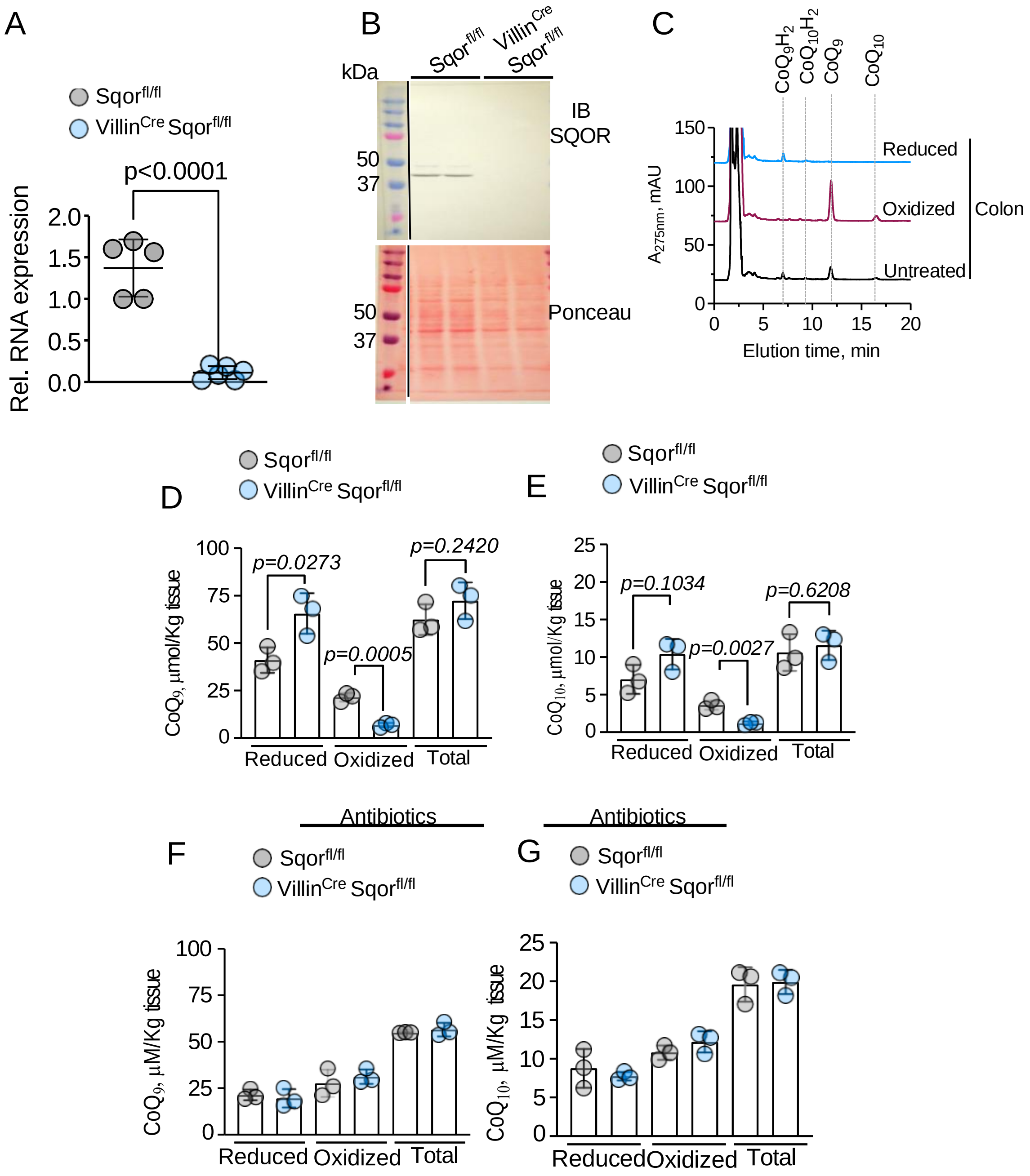


**Supplementary Figure 1. Validation of Sqor KO and CoQ quantitation. Related to Figure 1. (A,B)** Validation of Sqor knockout in colon through quantitative PCR (A), n=5 for Sqor$^{fl/fl}$ and n=6 for Villin$^{Cre}$ Sqor$^{fl/fl}$ and western blotting (B), n=2. (**C**) Colon lysate exhibit three peaks assigned as $CoQ_9H_2$, $CoQ_9$ and $CoQ_{10}$ (black); oxidation with *p*-benzoquinone (p-BQ) leads to the disappearance of the $CoQ_9H_2$ and an increase in the intensity of $CoQ_9$ and $CoQ_{10}$ peaks (red); reduction with sodium borohydride caused disappearance of $CoQ_9$ and $CoQ_{10}$ and increase in $CoQ_9H_2$ and $CoQ_{10}H_2$ peaks (blue). (**D,E**) $CoQ_9$ (E) and $CoQ_{10}$ pool sizes in Sqor$^{fl/fl}$ and Villin$^{Cre}$ Sqor$^{fl/fl}$ mice, n=3 independent mice. (**F,G**) $CoQ_9$ (G) and $CoQ_{10}$ (H) pool sizes in Sqor$^{fl/fl}$ and Villin$^{Cre}$ Sqor$^{fl/fl}$ mice on antibiotics, n=3 independent mice.

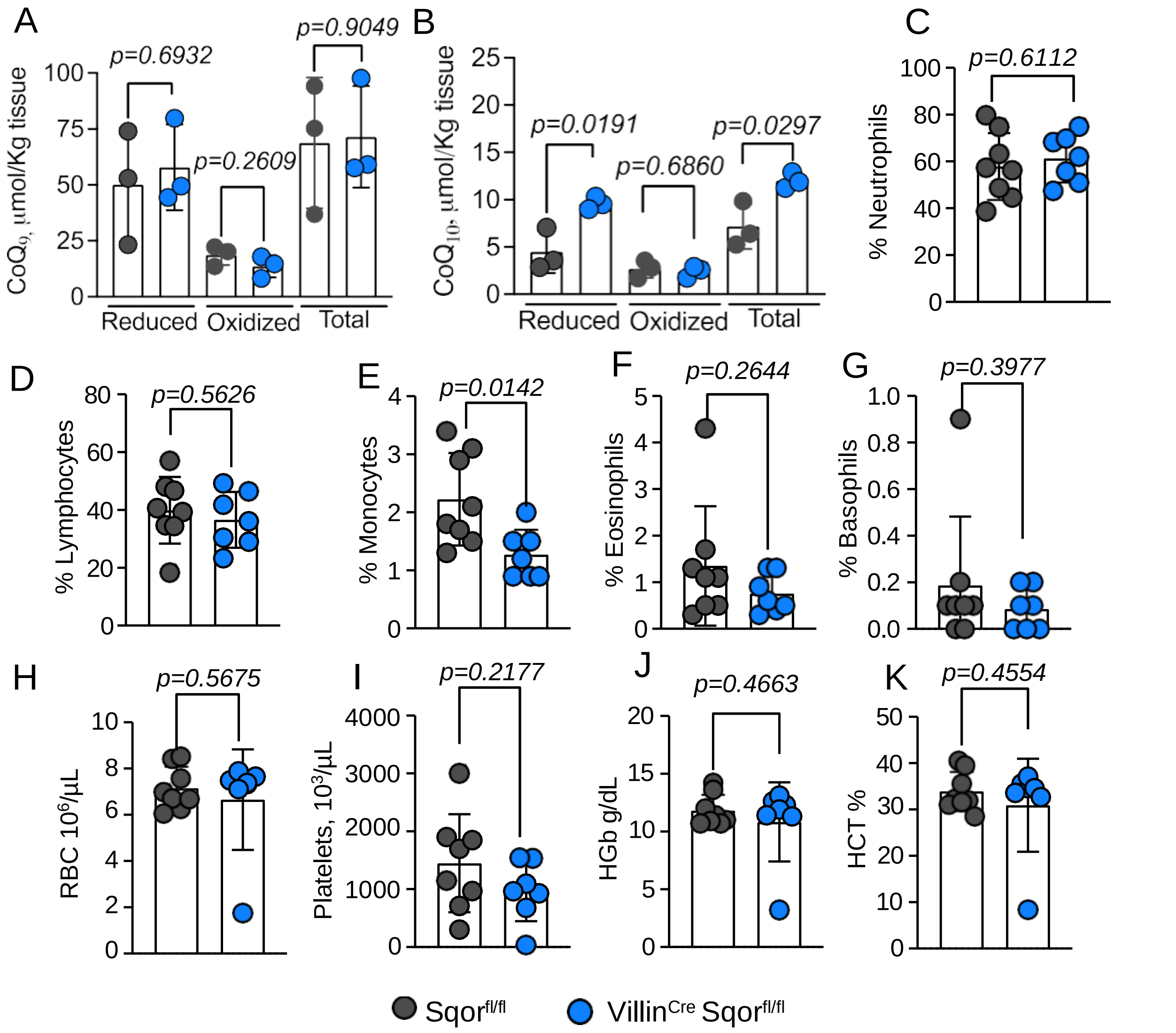


**Supplementary Figure 2. Complete blood count and CoQ analysis. Related to Figure 2**. (**A,B**) $CoQ_9$ (A) and $CoQ_{10}$ (B) pool sizes in $Sqor^{fl/fl}$ and $Villin^{Cre}$ $Sqor^{fl/fl}$ mice on 1.5% MRD. (**C-K**) Blood cell profiling of $Sqor^{fl/fl}$ and $Villin^{Cre}$ $Sqor^{fl/fl}$ mice on 1.5% MRD.

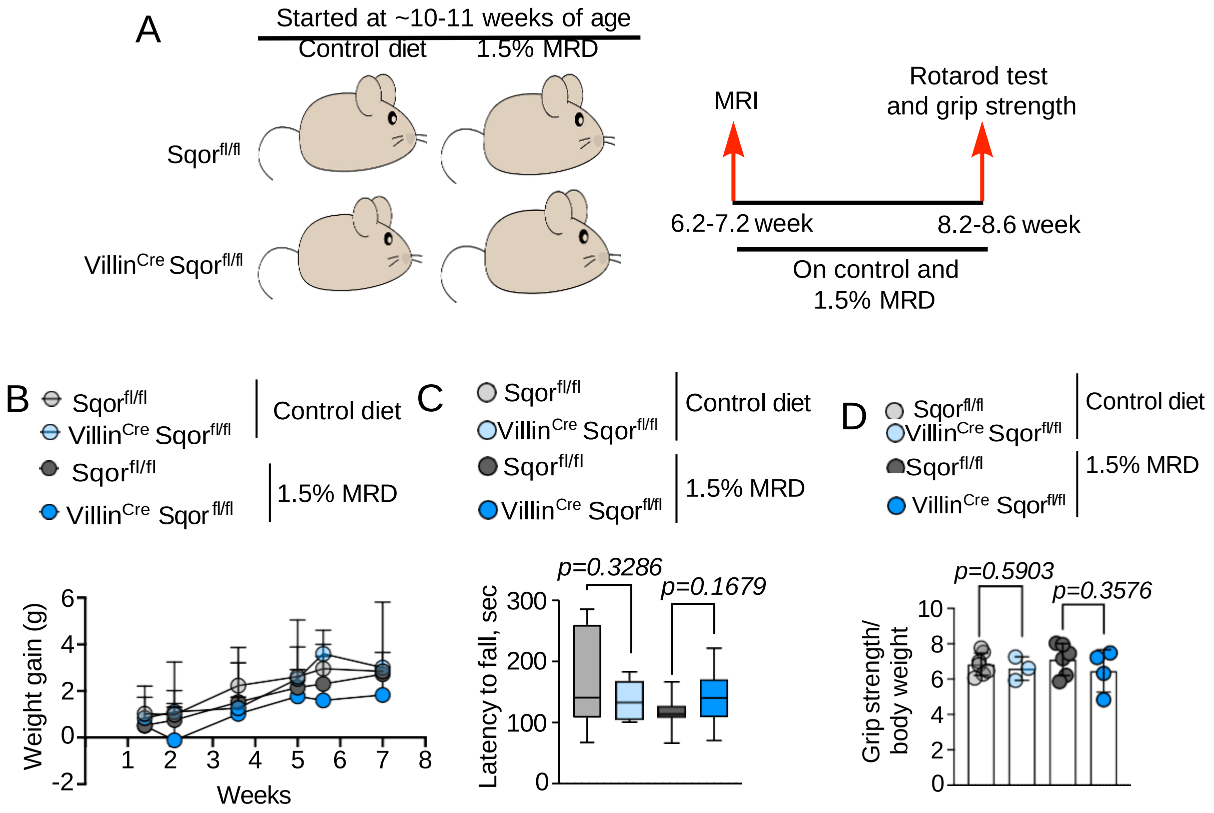


**Supplementary Figure 3**. **Related to Figure 3**. **Effect of MRD on Villin$^{Cre}$ Sqor$^{fl/fl}$ mice**. (**A**) Scheme describing the experimental setup. (**B**) Weight gain in Sqor$^{fl/fl}$ and Villin$^{Cre}$ Sqor$^{fl/fl}$ mice on control and 1.5% MRD. (**C**) Latency to fall in rotarod test to assess motor strength. (**D**) Comparison of grip strength between the genotypes and diets.

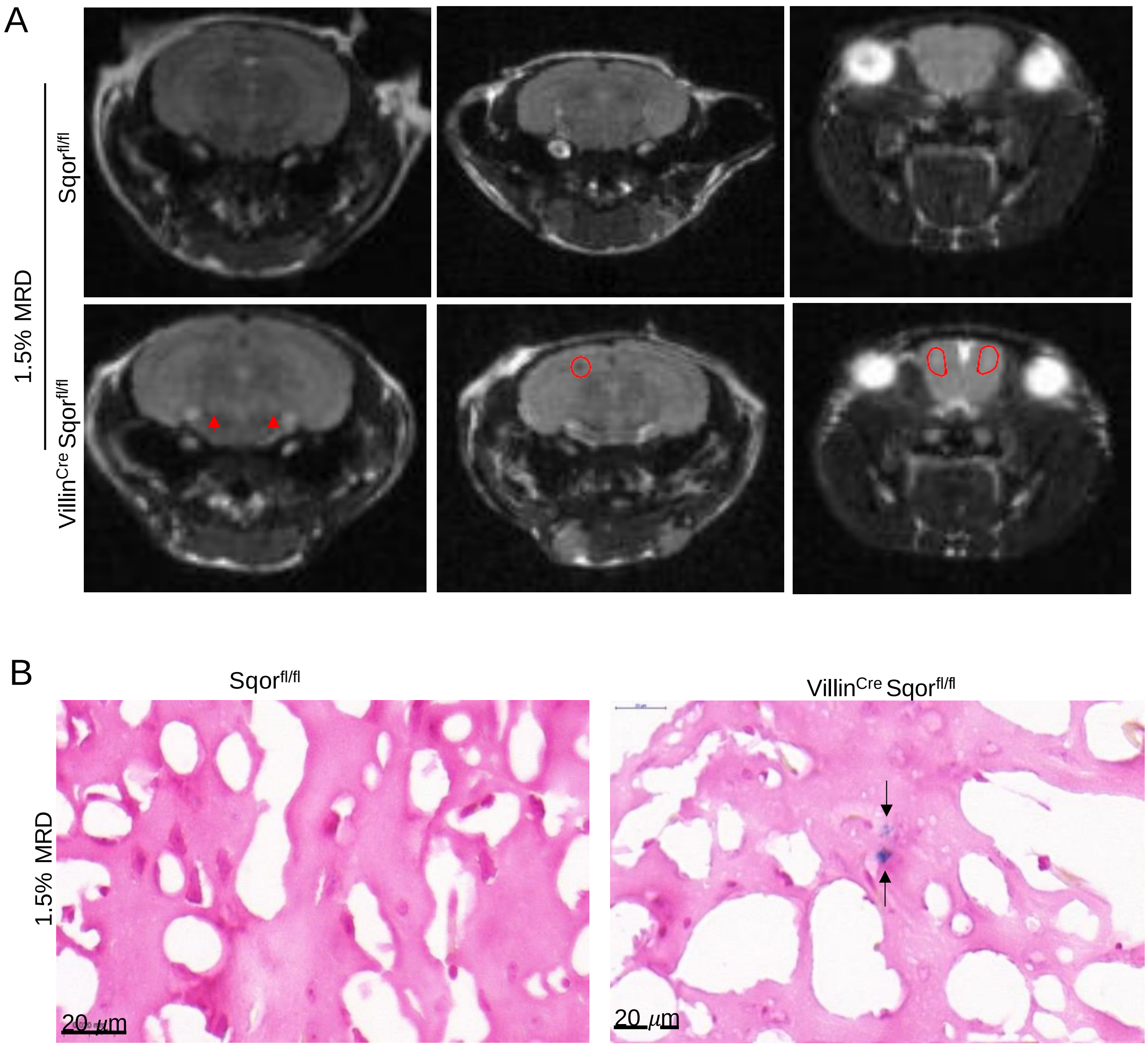


**Supplementary Figure 4. Related to Figure 3. Effect of MRD on brain MRI in Sqor$^{fl/fl}$ and Villin$^{Cre}$ Sqor$^{fl/fl}$ mice.** (**A**) T2 MRI showing hypointensity (circles) and hyperintensity (arrowheads) in Villin$^{Cre}$ Sqor$^{fl/fl}$ compared to Sqor$^{fl/fl}$ mice on a 1.5% MRD. (B) Prussian blue staining of brain sections showing free iron accumulation (black arrows) in Villin$^{Cre}$ Sqor$^{fl/fl}$ compared to Sqor$^{fl/fl}$ mice on a 1.5% MRD.

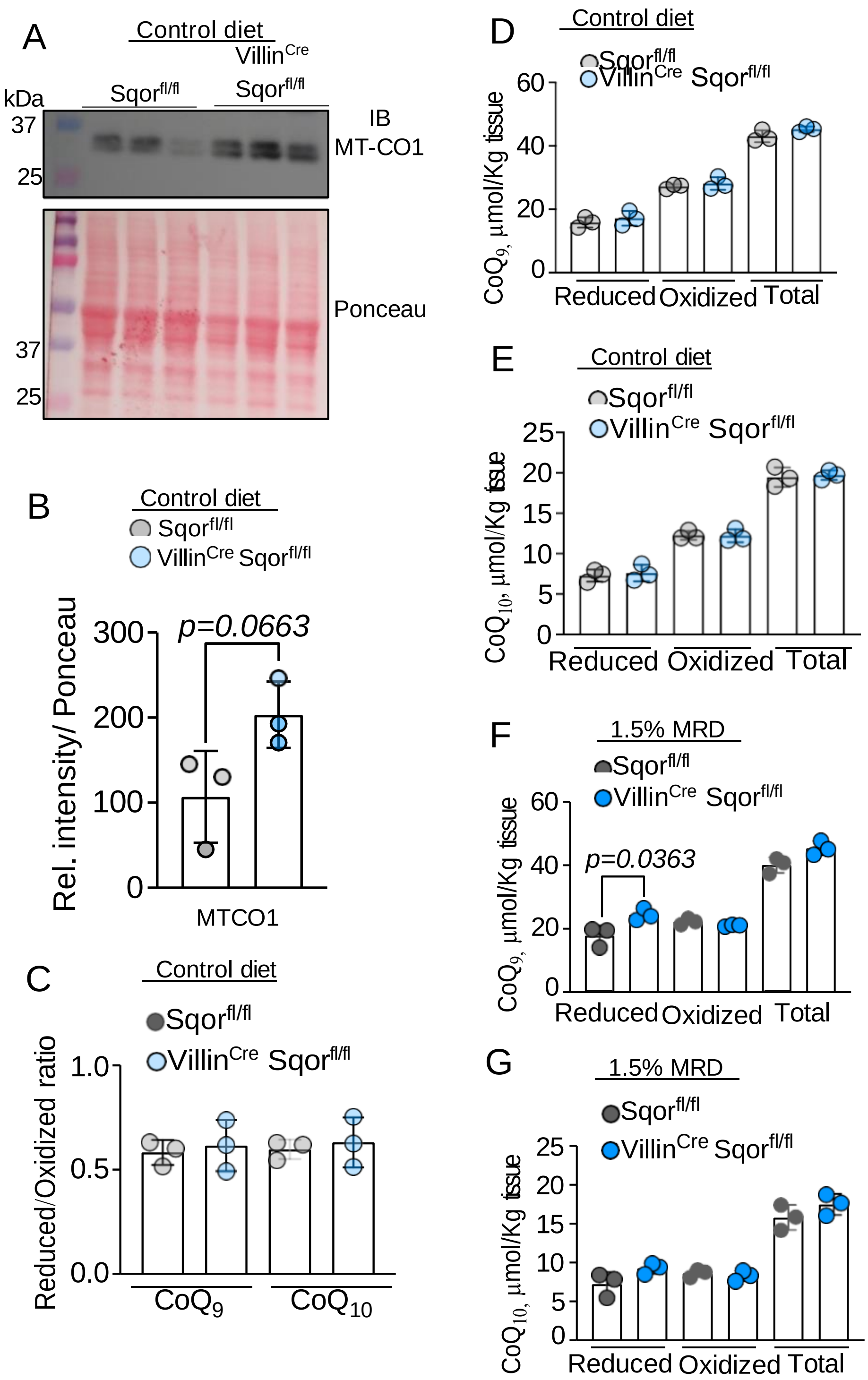


**Supplementary Figure 5. Related to Figure 3**. **Effect of MRD on brain bioenergetics in control versus Villin$^{Cre}$ Sqor$^{fl/fl}$ mice.** (**A,B**) Western blot (A) and quantitative analysis (B) of MT-CO1 levels in murine brain on a control diet (n= 3 independent mice). (**C-G**) Brain $CoQ_9$ and $CoQ_{10}$ redox (C) and pool sizes (D-G) in Sqor$^{fl/fl}$ and Villin$^{Cre}$ Sqor$^{fl/fl}$ mice on control versus 1.5% MRD.

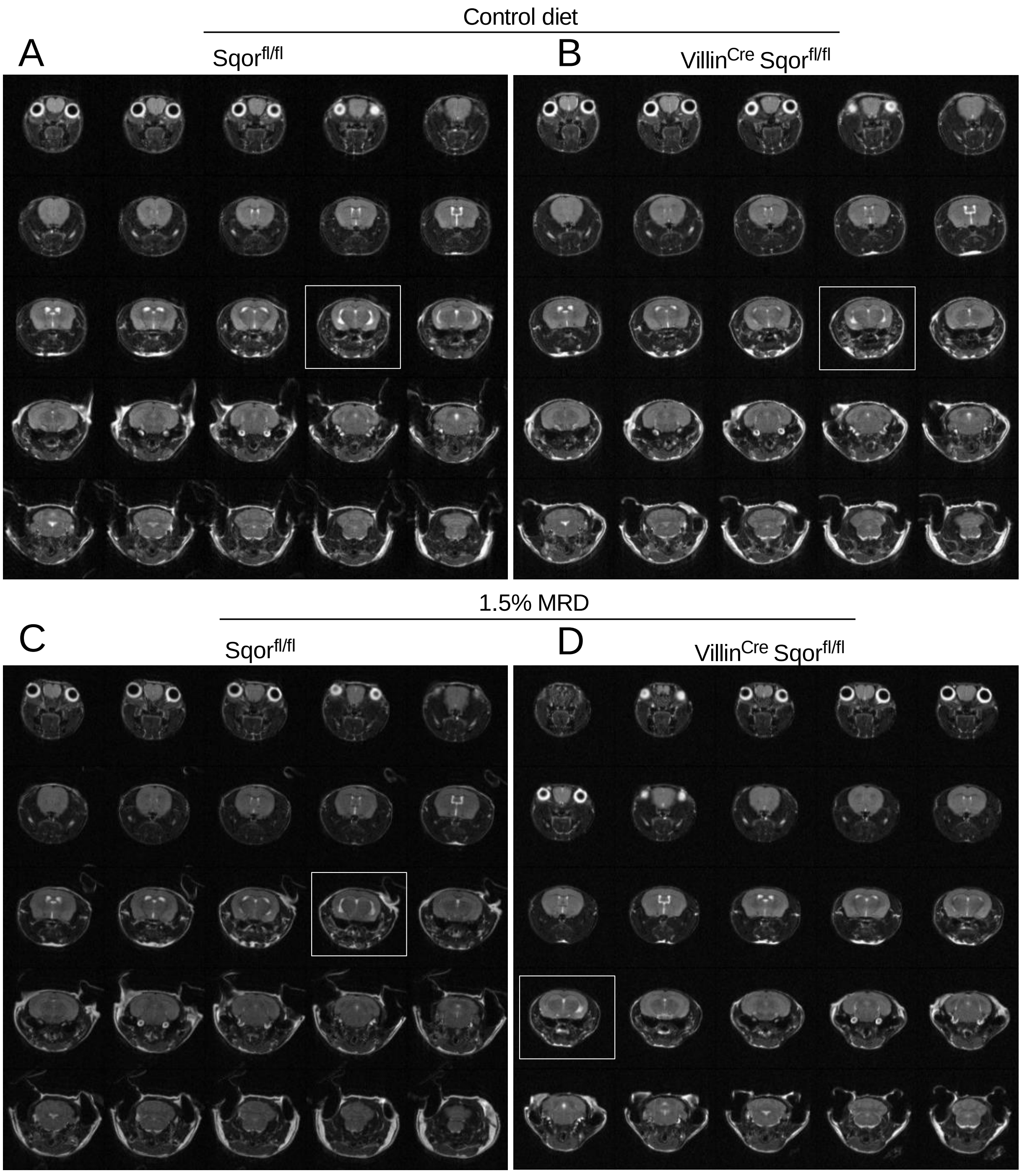


**Supplementary Figure 6. Related to Figure 3. Comparison of T2-weighted brain MRIs. A-D**. T2-weighted MRI of axial brain sections from $Sqor^{fl/fl}$ and $Villin^{Cre}$ $Sqor^{fl/fl}$ mice (**A,B**) on control and 1.5% MRD (**C,D**). The white outlines highlight sections in which asymmetric loss of lateral ventricles is evident in $Villin^{Cre}$ $Sqor^{fl/fl}$ mice on a 1.5% MRD compared to the other samples.

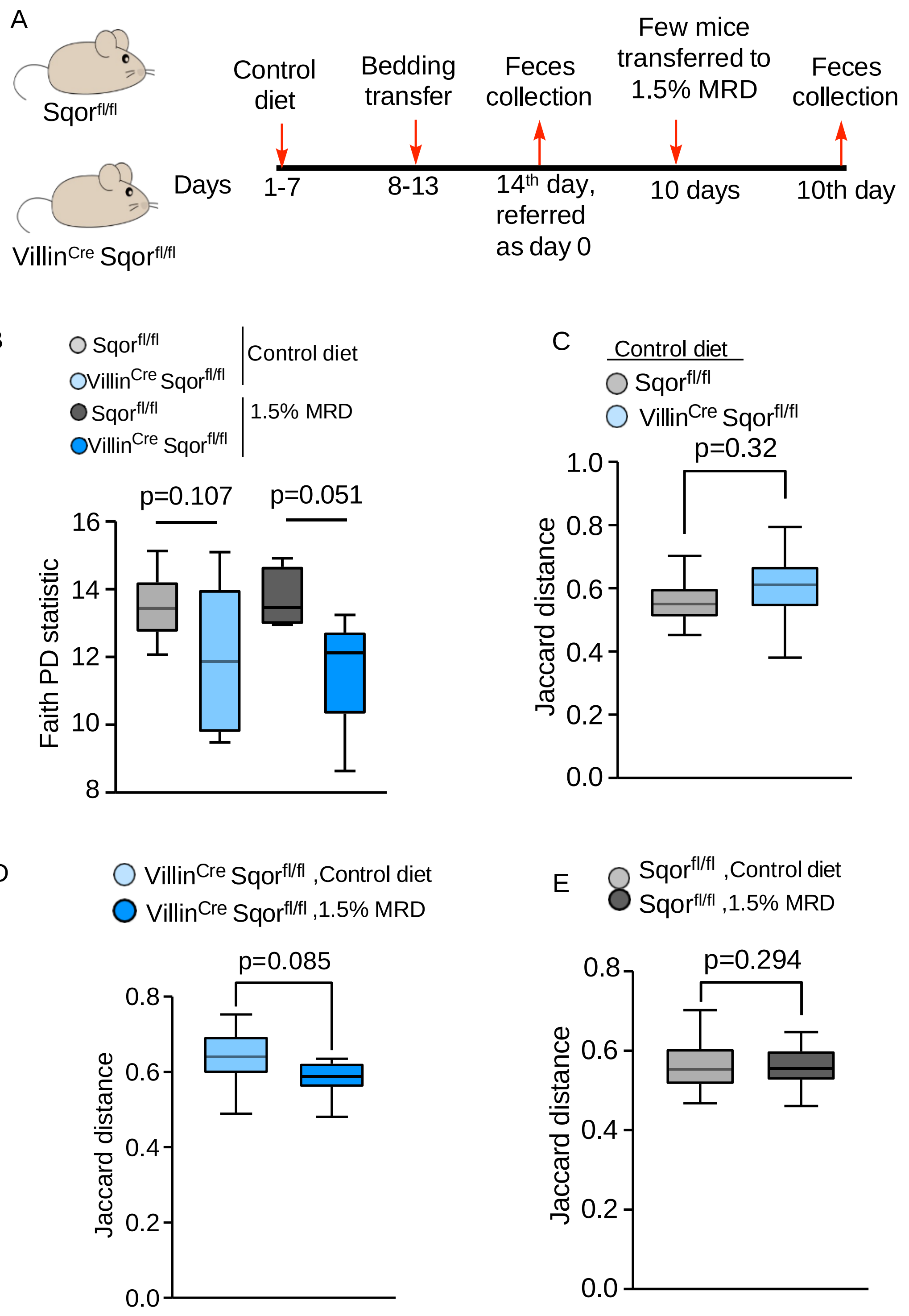


**Supplementary Figure 7. Related to Fig 4. Alpha diversity and beta diversity comparisons in control versus Villin$^{Cre}$ Sqor$^{fl/fl}$ mice on control or 1.5% MRD.** (**A**) Scheme showing mice fecal harvest for 16S rRNA analysis. (**B**) Alpha diversity as measured by Faith's PD statistic, with p values calculated by two-tailed t test. (**C-E**) Jaccard distance within and between group pairwise comparisons using PERMANOVA to calculate adjusted p values.
